# ASCL1 regulates neurodevelopmental transcription factors and cell cycle genes in glioblastoma

**DOI:** 10.1101/2020.02.20.958132

**Authors:** Tou Yia Vue, Rahul K. Kollipara, Mark D. Borromeo, Tyler Smith, Tomoyuki Mashimo, Dennis K. Burns, Robert M. Bachoo, Jane E. Johnson

## Abstract

Glioblastomas (GBMs) are incurable brain tumors with a high degree of cellular heterogeneity and genetic mutations. Transcription factors that normally regulate neural progenitors and glial development are aberrantly co-expressed in GBM, conferring cancer stem-like properties to drive tumor progression and therapeutic resistance. However, the functional role of individual transcription factors in GBMs *in vivo* remains elusive. Here, we demonstrate that the basic-helix-loop-helix (bHLH) transcription factor ASCL1 regulates transcriptional targets that are central to GBM development, including neural stem cell and glial transcription factors, oncogenic signaling molecules, chromatin modifying genes, and cell cycle and mitotic genes. We also show that the loss of ASCL1 significantly reduces the proliferation of GBMs induced in the brain of a genetically relevant glioma mouse model, resulting in extended survival times. RNA-seq analysis of mouse GBM tumors reveal that the loss of ASCL1 is associated with downregulation of cell cycle genes, illustrating an important role for ASCL1 in controlling the proliferation of GBM.

**TABLE OF CONTENTS:** *Main Points:*

- ASCL1 is co-expressed with neural stem cell/glial transcription factors in GBM
- ASCL1 binds to genes that are important for cell proliferation and cancer in the brain.
- Loss of ASCL1 downregulates cell cycle genes and increase survival of glioma mouse model.

*Table of Content Image:* 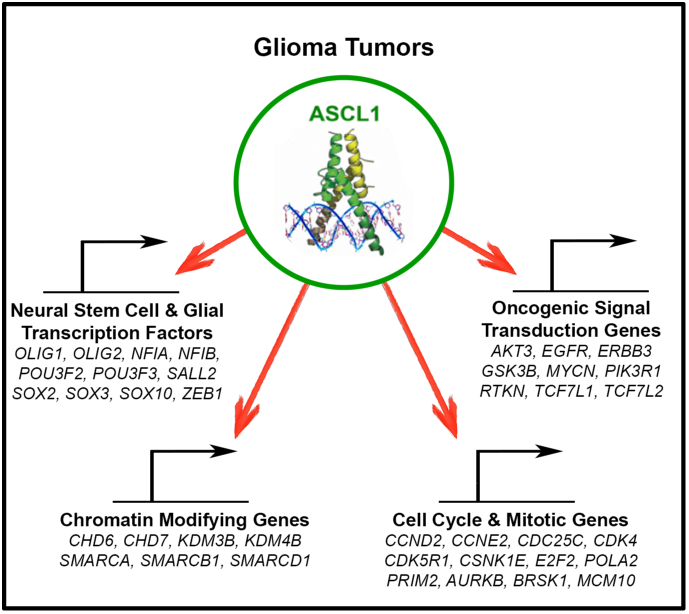

## INTRODUCTION

Glioblastomas (GBMs) are incurable brain tumors most commonly found in adults. Despite significant advances in imaging and surgical resection techniques combined with aggressive radiotherapy and chemotherapy, the median survival for GBM patients remains stagnated between 14-16 months, with greater than 90% of patients succumbing to their disease within 5 years of diagnosis (Ostrom et al., 2016). A major reason for this poor prognosis is due to the high degree of heterogeneity and plasticity of these neoplasms, and the lack of mechanistic insights into the pan-therapeutic resistance of GBM tumor cells (Babu et al., 2016; Brennan et al., 2013; Lathia, Heddleston, Venere, & Rich, 2011b).

Concerted sequencing efforts from the Cancer Genome Atlas (TCGA) Research Network revealed a complex somatic landscape for GBMs involving oncogenes (*BRAF, EGFR, PDGFRα, MET, PIK3C, MYCN*), tumor suppressor genes (*CDNK2A/B, PTEN, NF1, RB1*) and chromatin modifying genes, which converge to activate signaling pathways (pAKT, Ras/MAPK, STAT) to promote tumor proliferation and growth (2008; Brennan et al., 2009; Brennan et al., 2013; Verhaak et al., 2010). Emerging evidence also suggests that a cellular hierarchy may exist within the heterogeneous GBM tumor composition, where a subpopulation of quiescent cancer stem-like cells, or glioma-stem-cells (GSCs), are postulated to be responsible for driving tumor growth, progression, and the development of resistance to therapeutic treatments (Bao et al., 2006; Chen et al., 2012; Lan et al., 2017; Lathia et al., 2011b; Lathia et al., 2011a; Parada, Dirks, & Wechsler-Reya, 2017).

Despite displaying an aberrant array of mutations, GSCs are universally marked by co-expression of a combination of transcription factors, some of which include ASCL1, NFIA, NKX2.2, OLIG2, POU3F2, SALL2, SOX2, and ZEB1 (Glasgow et al., 2017; Lu et al., 2016; Rheinbay et al., 2013; Singh et al., 2017; Suva et al., 2014). These transcription factors have been extensively studied in the developing central nervous system (CNS), where each has been shown to regulate the fate, proliferation and/or migration of neural progenitor and glial precursor cells in stage specific processes. In the context of gliomas, these transcription factors are often constitutively co-expressed and have been shown to function in a combinatorial manner in determining the tumorigenicity and differentiation status of tumor cells (Gangemi et al., 2009; Ligon et al., 2007; Rheinbay et al., 2013; Singh et al., 2017; Suva et al., 2014).

In this study, we focus on ASCL1, a class II basic-helix-loop-helix (bHLH) transcription factor that forms a heterodimer with class I bHLH E-proteins (such as E47/TCF3) to activate specific target genes (Kageyama, Ohtsuka, Hatakeyama, & Ohsawa, 2005). During embryogenesis, ASCL1 is expressed in specific populations of neural progenitor domains and glial precursor cells throughout the neural tube from the spinal cord to the brain (Helms et al., 2005; Parras et al., 2004; Parras et al., 2007; Sugimori et al., 2007; Sugimori et al., 2008; Vue, Kim, Parras, Guillemot, & Johnson, 2014), including in neurogenic regions of the adult brain (Kim, Leung, Reed, & Johnson, 2007; Kim, Ables, Dickel, Eisch, & Johnson, 2011). Recently, ASCL1 was shown to be capable of reorganizing and promoting the accessibility of closed chromatin in embryonic stem cells as well as glioma cell lines (Casey, Kollipara, Pozo, & Johnson, 2018; Raposo et al., 2015). Not surprisingly, genome wide profiling revealed a critical role for ASCL1 in interacting with both Wnt and Notch signaling pathways to control the tumorigenicity of glioma cells in culture (Park et al., 2017; Rheinbay et al., 2013). To date however, whether ASCL1 is absolutely required for glioma tumor development in the brain as it has been shown for a mouse model of small-cell-lung-carcinoma (SCLC) (Borromeo et al., 2016) remains to be determined. Here, we sought to identify the direct *in vivo* role and transcriptional targets of ASCL1 in brain tumors of previously characterized patient-derived-xenograft (PDX)-GBM and genetically engineered glioma mouse models.

## MATERIALS AND METHODS

### Glioma Mouse Models

Patient-derived-xenograft (PDX) GBM (R738 and R548) were passaged orthotopically in the brains of NOD-SCID mice as previously described (Marian et al., 2010; Marin-Valencia et al., 2012). Generation and genotyping of mouse strains used to generate the glioma models were as previously reported: *Glast*^*CreERT2*^ knock-in (Mori et al., 2006); *Ascl1*^*GFP*^ knock-in [Ascl1^tm1Reed^/J 012881] (Kim et al., 2007); *Ascl1*^*F*^ [Ascl1-floxed] (Andersen et al., 2014; Pacary et al., 2011); *Nf1*^*F*^ [Nf1^tm1Par^/J 017639] (Zhu et al., 2001); *p53*^*F*^ [p53-floxed] (Lin et al., 2004); and the Cre reporter lines *R26R*^*LSL-YFP*^ [Gt(ROSA)26Sor^tm1(EYFP)Cos^/J 006148] (Srinivas et al., 2001) and *R26R*^*LSL-tdTOM*^ [Gt(ROSA)26Sortm^14(CAG-tdTomato)Hze^/J 013731] (Madisen et al., 2010). All animal procedures followed NIH guidelines and were approved by the UT Southwestern Institutional Animal Care and Use Committee.

### Mouse Breeding and Tamoxifen Administration

The appearance of a vaginal plug was considered embryonic day (E) 0.5 and the day of birth was noted as postnatal day (P)0. To induce tumor formation in the brains of *Glast*^*CreERT2/+*^*;Nf1*^*F/F*^*;Trp53*^*F/F*^ mice, tamoxifen (Sigma T5648, dissolved in 10% ethanol/90% sunflower oil) was administered intraperitoneally (62.5 mg/kg body weight) to pregnant females at E14.5. Due to the effects of tamoxifen on birth complications, cesarean section was performed and pups were carefully introduced and raised by a foster female.

### Tissue preparation, H&E staining, and Immunofluorescence

Tumor bearing mice were trans-cardiac-perfused with 4% PFA in PBS. Brains were submerged in 30% sucrose/PBS at 4°C, and embedded in O.C.T. for cryosectioning. H&E staining of tumors was done by the UT Southwestern Histopathology Core. Grading of brain tumors was determined by a board certified neuropathologist.

For immunohistochemistry, tissue sections were incubated with primary antibody in 1% goat or donkey serum/0.3% Triton X-100/PBS overnight, followed by incubation with secondary antibody conjugated with Alexa Fluor 488, 568 or 647 (Molecular Probes), and coverslipped with Vectashield (#101098-042) for confocal microscopy (LSM 510 & 720). The following antibodies were used:

| Primary Antibodies | Source & Catalogue Number | Dilution |
| --- | --- | --- |
| Chicken Anti-GFP | Chemicon, AB16901 | 1:500 |
| Goat Anti-SOX10 | R&D Systems, AF2864 | 1:20 |
| Guinea Pig Anti-ASCL1 | Kim et al., 2008, TX518 | 1:1,000 – 1:10,000 |
| Mouse Anti-GFAP | Sigma, G3893 | 1:500 |
| Mouse Anti-MBP | Calbiochem, NE1019 | 1:300 |
| Mouse Anti-NEUN | Chemicon, MAB377 | 1:1,000 |
| Rabbit Anti-Ki67 | Abcam, ab15580 | 1:500 |
| Rabbit Anti-OLIG2 | Millipore, AB9610 | 1:1,000 |
| Rabbit Anti-SOX2 | Millipore, AB5603 | 1:1,000 |
| Rat Anti-PDGFRα (APA5) | BD Pharmingen, 558774 | 1:100 |

### ChIP-seq, RNA-seq, and Data Analysis

Two independent ASCL1 ChIP-seq experiments were performed using PDX-GBMs (R548 and R738) dissected from brains of NOD-SCID mice exhibiting symptoms of the presence of tumor. Briefly, as previously described (Borromeo et al., 2016), tumor tissues were homogenized and fixed in 1% formaldehyde to crosslink proteins and DNA, followed by quenching with 0.125 M of glycine. Nuclear chromatin was pelleted, washed with cold PBS, and sonicated into 200-300bp fragments using a Biorupter (Diagenode). A 10% portion of the sheared chromatin was set aside as input DNA. Approximately 100μg was subjected to immunoprecipitation using ∼5 μg of mouse anti-ASCL1 (Mash1) antibody (BD Biosciences, 556604). Washes and reverse-crosslinking were performed using Dynabeads Protein G to elute ChIP DNA.

For RNA-seq experiments, the brain tumors were carefully dissected to enrich for tumor tissues and total RNA was extracted using a Direct-zol RNA MiniPrep Kit (Zymo Research). RNA integrity number (RIN) for all tumors was determined to be between 8-10 using a Bioagilent Analyzer. ChIP DNA and input DNA from PDX-GBMs and total RNAs from mouse brain tumors were sent for library preparation and sequencing on an Illumina High-Seq 2000 at the UT Southwestern Next Generation Sequencing Core.

To analyze ASCL1 ChIP-seq data, sequence reads were aligned to the human reference genome (hg19) using bowtie2 (v.2.2.6) (Langmead & Salzberg, 2012). Low-quality reads and duplicate reads were removed from aligned files using “samtools view -bh-F 0 × 04 -q 10” (v1.2) (Li, 2011) and “Picard MarkDuplicates.jar” (v. 1.131) commands (Picard 2018, Broad Institute, GitHub repository). The ChIP-seq signal enriched regions were identified using the “findPeaks” module available in HOMER software (v.4.7) (Heinz et al., 2010). The ChIP-seq signal shown in UCSC browser tracks are normalized read counts. *De novo* motif discovery and analysis were performed using “findMotifsGenome” module available in HOMER software (v.4.7). A 150 bp region around the peak summit was used to identify the primary binding motif and other potential DNA-binding co-Lanfactor motifs.

To analyze mouse tumor RNA-seq data, sequenced reads were aligned to the mouse mm10 genome using TopHat 2.1.0 (Kim et al., 2013). Default settings were used, with the exception of –G, specifying assembly to the mm10 genome, --library-type fr -first strand, and – no-novel-juncs, which disregards noncanonical splice junctions when defining alignments. DESeq2 (Love, Huber, & Anders, 2014) was used to incorporate RNA-seq data from the five biological replicates for *Ascl1*^*WT*^ and *Ascl1*^*CKO*^ tumor samples, and differentially expressed genes were identified using default parameters.

To investigate the similarity/difference between *Ascl1*^*WT*^ and *Ascl1*^*CKO*^ tumors in comparison to each other and to CNS cell types, multidimensional scaling (MDS) was performed using the plotMDS function available in edgeR package (Robinson, McCarthy, & Smyth, 2010). Finally, to identify enrichment of gene signature sets in rank ordered gene lists obtained from *Ascl1*^*WT*^ and *Ascl1*^*CKO*^ tumor samples, gene set enrichment analysis (GSEA) (Subramanian et al., 2005) was performed and the signal-to-noise ratio metric was used to rank the genes.

### GBM Subtype Classification and Heatmap Clustering Analyses

The GBM subtype signatures defined by Verhaak et al. (Verhaak et al., 2010) were used for hierarchical clustering for 164 GBM patient samples and 5 normal brains from TCGA for which RNA-seq data was available (2008; Brennan et al., 2009; Brennan et al., 2013; Verhaak et al., 2010). Spearman rank order correlation and ward.D2 clustering method were applied to identify the various GBM subtypes. Heatmaps were generated using absolute expression values (RPKM) for the selected list of genes or significantly changed genes, and hierarchical clustering was performed using the correlation distance metric and the ward.D2 method using the heatmap.2 function available in the *gplots* R package.

### Gene Targets and Pathway Enrichment Analysis

To identify ASCL1 putative targets, genes associated with the ASCL1 ChIP-seq peaks were annotated using GREAT v3.0.0 (http://great.stanford.edu/public/html/) (McLean et al., 2010), which was then cross-referenced with the top 10% of genes (2,136) whose expression positively correlates with ASCL1 expression by computing the Spearman rank order correlation (>0.4) using RNA-seq of TCGA GBM expression data. An overlap of 1,106 genes was identified as ASCL1 putative target genes. These genes were then subjected to pathway enrichment analysis performed using ConsensusPathDB (http://cpdb.molgen.mpg.de/) (Herwig, Hardt, Lienhard, & Kamburov, 2016). Relevant significantly enriched overrepresented gene sets (FDR ≤ 5%) were selected for illustration.

### Quantification of ASCL1+, OLIG2+, SOX2+ and Ki67+ Tumor Cells

The number of DAPI+ tumor cells that were ASCL1+ along with each of the various markers were quantified using Image J on 20X immunofluorescence confocal images of both R548 and R738 PDX-GBMs. Quantifications were performed on at least three images taken from different areas per tumor for each marker (N=4).

To determine the expression of ASCL1, OLIG2, and SOX2 in human GBMs, RNA-seq of 164 TCGA primary GBM and 5 normal brain samples were analyzed and categorized into the various subtypes using the 840 GBM Subtype Signature Genes (Verhaak et al., 2010). Average RPKM for *ASCL1, OLIG2*, and *SOX2* was determined for each GBM subtype. Outlier samples exhibiting an RPKM value > 2 standard deviations away from the mean were excluded.

To compare the Ki67 index between *Ascl1*^*WT*^ (N=6) or *Ascl1*^*CKO*^ (N=5) tumors, 20X immunofluorescence confocal images were taken from three different areas per tumor. Because the distribution of Ki67+ cells is not uniform within a large growing tumor, we limited our imaging to only regions with the highest density of Ki67+ cells. Quantification of the number of Ki67+;DAPI+/total DAPI+ cells was then performed blind of genotype for each image and compiled for comparison between *Ascl1*^*WT*^ or *Ascl1*^*CKO*^ tumors using a Wilcox test.

## RESULTS

### Neurodevelopmental transcription factors ASCL1, OLIG2, and SOX2 are highly co-expressed in human GBMs

ASCL1, OLIG2, and SOX2 have previously been reported to be expressed in GBMs (Gangemi et al., 2009; Ligon et al., 2007; Lu et al., 2016; Park et al., 2017; Rheinbay et al., 2013; Singh et al., 2017; Somasundaram et al., 2005). However, the extent to which these factors are co-expressed in GBM tumors *in vivo* remains unclear. Using patient-derived-xenograft (PDX)-GBM lines (R548, R738), in which tumors from patients were passaged orthotopically in the brains of NOD-SCID mice (Figure 1A) (Marian et al., 2010; Marin-Valencia et al., 2012), we demonstrated that the transplanted tumors exhibit pathological characteristics of gliomas (Figure 1B, C) and express ASCL1, OLIG2, and SOX2 in the majority of tumor cells (Figure 1D-M). Quantification shows that each transcription factor occupied 74%, 81%, and 85% of tumor cells counterstained with DAPI, respectively (Figure 1N). Co-localization analysis revealed that 48% of ASCL1+ tumor cells were positive for the proliferation marker Ki67 (Figure 1K-M, O), whereas over 90% of ASCL1+ cells were OLIG2+ and SOX2+ (Figure 1O), indicating that these three transcription factors are co-expressed in the majority of the PDX-GBM cells *in vivo*.

**Figure 1.**
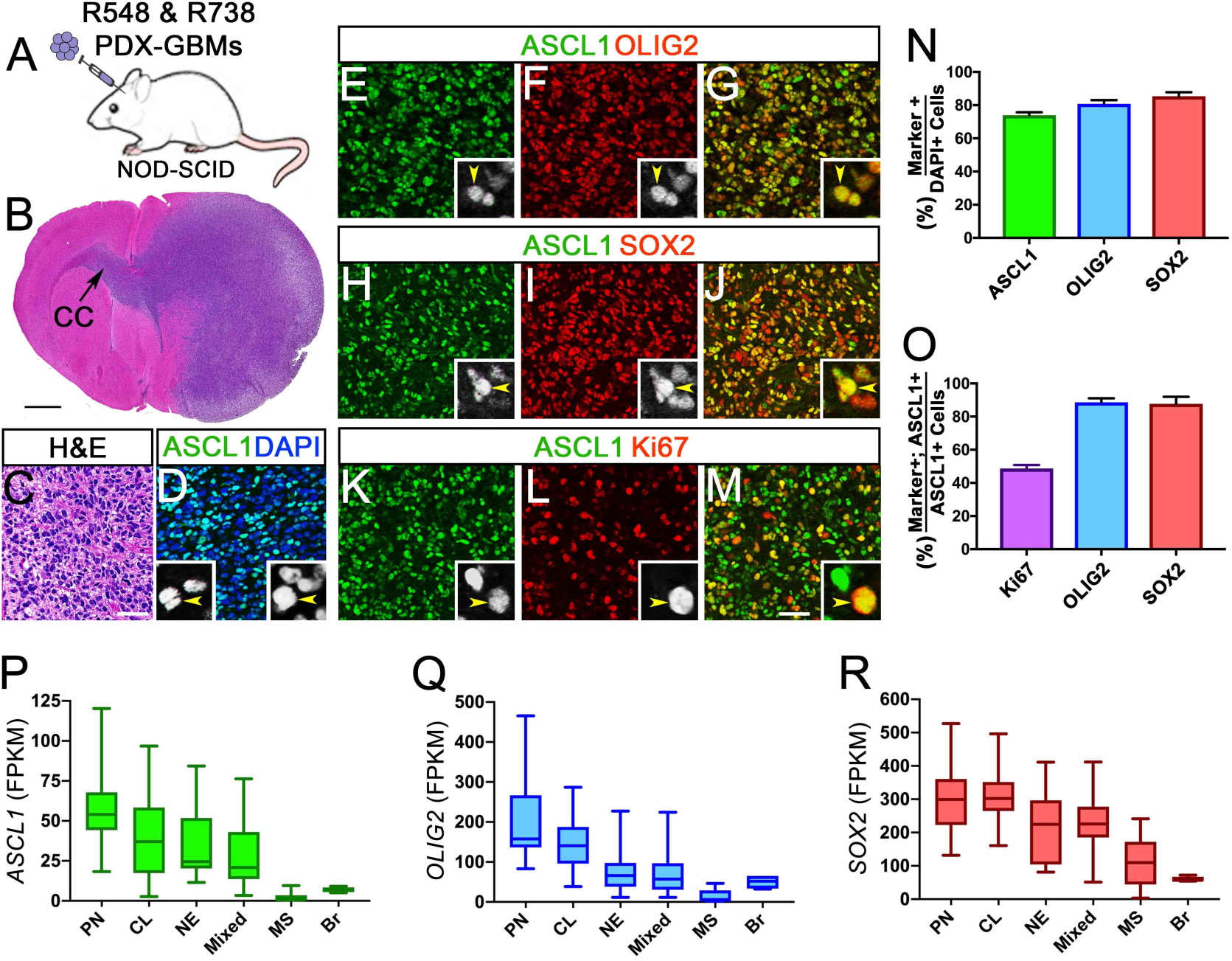
Neurodevelopmental transcription factors ASCL1, OLIG2, and SOX2 are highly expressed in the majority of GBMs. **(A)** Schematic of PDX-GBMs (R548 and R738) grown orthotopically in the brains of NOD-SCID mice. **(B, C)** H&E staining showing tumor is a high-grade glioma and is migrating across the corpus callosum (CC). **(D-M)** Immunofluorescence showing co-expression of ASCL1 with OLIG2 (E-G), SOX2 (H-J), and Ki67 (K-M) in the PDX-GBMs. **(N, O)** Quantification of the percentage of DAPI+ tumor cells that are ASCL1+, OLIG2+, or SOX2+ (N), and the percentage of ASCL1+ tumor cells that are also Ki67+, OLIG2+, or SOX2+ (O). N=4 PDX-GBM. **(P-R)** Box whisker plot of RNA-seq data from 160 TCGA Primary GBMs and 5 normal brain samples (Brennan et al, 2013) demonstrating that *ASCL1* (P), *OLIG2* (Q), and *SOX2* (R) are highly expressed in the majority of GBM subtypes but are low in MS subtype and normal brain (Br). GBM subtype was determined using the 840 GBM Subtype Signature Genes (Verhaak et al, 2010). PN-proneural, MS-mesenchymal, CL-classical, NE-neural. Mixed GBM subtype express multiple subtype signatures. Scale bar is 1 mm for B and 50 μm for C-M, and 12.5 μm for all insets in D-M.

We next sought to determine the expression level of ASCL1, OLIG2, and SOX2 across primary GBMs exhibiting a variety of genomic alterations. Leveraging RNA-seq data of 164 TCGA primary GBMs, along with 5 normal control brain samples (Brennan et al., 2013), we first classified these primary GBMs into the four GBM subtypes (proneural, neural, classical, mesenchymal) as previously defined using an 840 gene list (Verhaak et al., 2010). Notably, while 107 samples can be classified into one of the four GBM subtypes, the remaining 57 samples expressed signatures of more than one subtype, which we referred collectively to as mixed GBMs (Figure S1). This finding echoes previous reports demonstrating the presence of multiple GBM subtype identities in different regions or cells of the same GBM tumors (Patel et al., 2014; Sottoriva et al., 2013). Expression of *ASCL1, OLIG2*, and *SOX2* across these GBM subtypes showed that they were highest in the proneural and classical subtypes, intermediate in the neural and mixed subtypes, but were extremely low in the mesenchymal subtype, even in comparison to normal brain (Figure 1P-R). Collectively, these findings illustrate that *ASCL1, OLIG2*, and *SOX2* are co-expressed at relatively high levels in the majority of primary GBMs with the exception of the mesenchymal subtype.

### ASCL1 binds to genes encoding neurodevelopmental and glial transcription factors, oncogene signaling molecules, and factors involved in cell cycle control and chromatin organization

Chromatin immunoprecipitation followed by deep sequencing (ChIP-seq) has previously been performed for ASCL1 in glioma cell lines in culture, and a dual role for ASCL1 was proposed to either promote or attenuate tumorigenicity depending on context (Park et al., 2017; Rheinbay et al., 2013). We performed ChIP-seq for ASCL1 in the two PDX-GBMs lines, both of which express high levels of ASCL1 (Figure 1), to identify its target genes *in vivo*. Using stringent peak calling criteria (Borromeo et al., 2014; Borromeo et al., 2016), we identified 9,816 statistically significant peaks in the genome of R548-PDX-GBM and 7,848 peaks in R738-PDX-GBM (blue rectangles, Figure 2A). Although only 4,207 of the significant peaks called overlapped in both PDX-GBMs, heatmaps of the ASCL1 ChIP-seq signal intensity, even for the non-significant peaks for each PDX-GBM, was noticeably higher than background for the combined 13,457 peaks called, indicating that the ASCL1 binding profile was similar in both PDX-GBMs (Figure 2A, Table S1).

**Figure 2.**
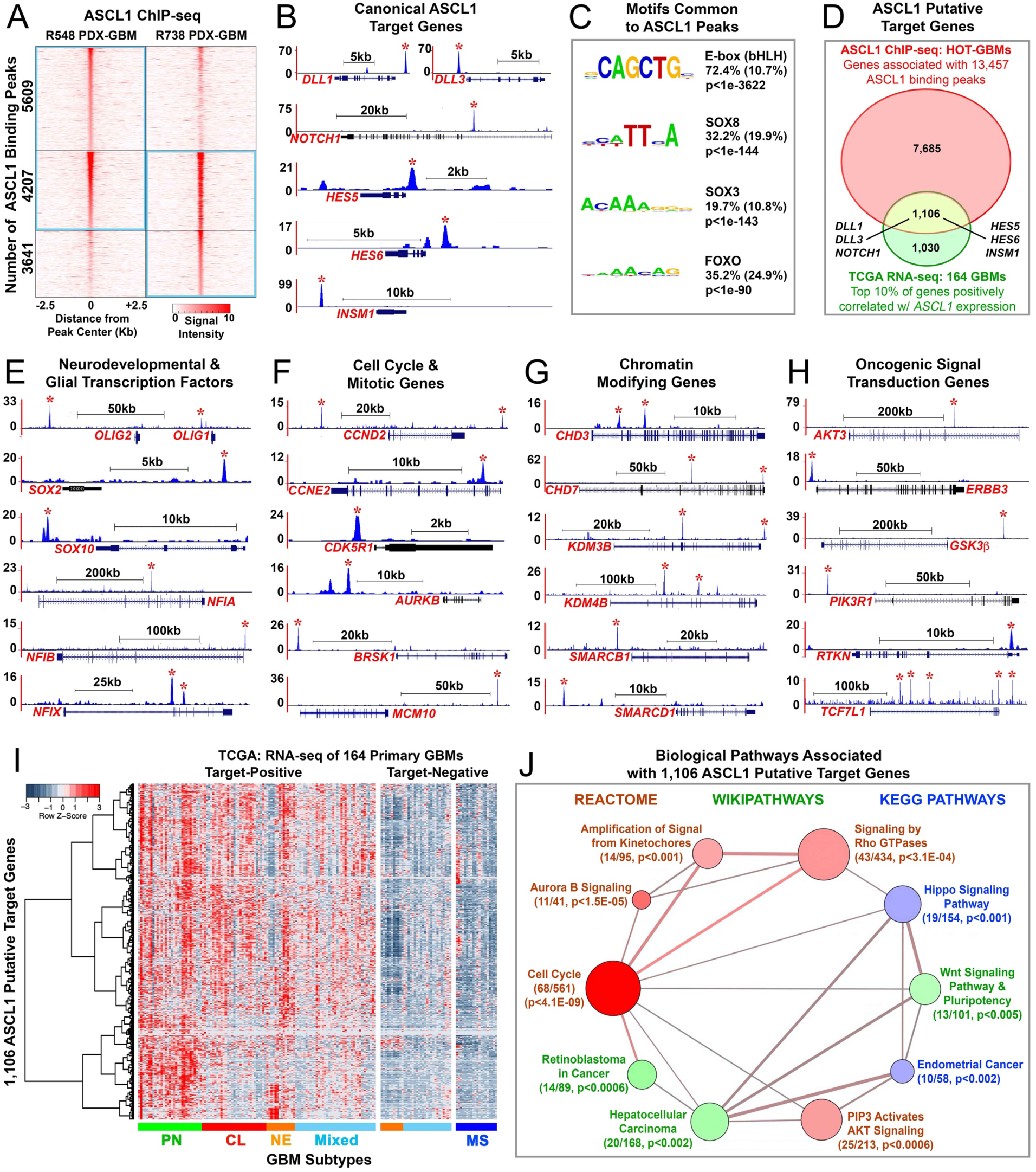
ASCL1 binds to target genes in GBMs involved in glial development, cell cycle progression, and cancer. **(A)** Heatmap of ASCL1 ChIP-seq signal intensity ±2.5 kb around 13,457 combined peaks identified in the genome of the PDX-GBMs. Blue rectangles indicate statistically significant peaks called by Homer. See Supplemental Table S1 for genomic coordinates of the ASCL1 binding sites. **(B)** ChIP-seq tracks of genomic regions surrounding canonical ASCL1 target genes *DLL1, DLL3, NOTCH1, HES5, HES6*, and *INSM1*. Asterisks indicate ASCL1 binding peaks meeting statistical criteria. **(C)** *De novo* motif analysis shows enrichment of bHLH E-box, SOX, and FOXO motifs directly beneath ASCL1 binding peaks. **(D)** Venn diagram intersecting genes (8,791, red oval) associated with ASCL1 binding peaks in the PDX-GBMs with the top 10% of genes (2,136, green oval) positively correlated (Spearmann corr<0.4) with *ASCL1* expression using RNA-seq data of 164 TCGA GBM samples (Supplemental Table S2). The overlap of 1,106 genes (yellow area) defines ASCL1 target genes, which included all the canonical ASCL1 target genes. **(E-H)** ChIP-seq tracks of ASCL1 binding peaks at loci of neurodevelopmental and glial transcription factors (E), cell cycle & mitotic genes (F), chromatin modifying genes (G), and oncogenic signaling pathway genes (H). **(I)** Heatmap and dendrogram illustrating relative expression of 1,106 ASCL1 putative target genes in GBM subtypes using RNA-seq of 164 TCGA primary GBM samples (Brennan et al, 2013). Note that ASCL1 target-positive GBMs include all subtypes except mesenchymal, while ASCL1 target-negative GBMs include all mesenchymal and some neural and mixed GBM subtypes. **(J)** Gene set over-representation analysis of 1,106 ASCL1 putative-target genes using ConsensusPathDB (cpdb.molgen.mpg.de). Biologically relevant enriched pathways are illustrated. Size of circle indicates the number of genes per pathway, size of edge indicates degree of gene overlaps between the pathways, and color indicates database sources. The number of ASCL1 putative-target genes over-represented in each pathway, and respective p-value are indicated. See Supplement Table S3 for complete gene set over-representation analysis.

To validate the quality and efficiency of our ChIP-seq, we next analyzed ASCL1 binding at known canonical targets (*DLL1, DLL3, NOTCH1, HES5, HES6* and *INSM1*) which have previously been shown to be directly regulated by ASCL1 in numerous contexts (Borromeo et al., 2014; Borromeo et al., 2016; Castro et al., 2011; Jacob et al., 2009; Park et al., 2017; Somasundaram et al., 2005; Ueno et al., 2012; Vias et al., 2008). As expected, ChIP-seq tracks revealed the presence of strong ASCL1 binding peaks at loci of all the canonical target genes examined (asterisk, Figure 2B). Moreover, ASCL1 is known to preferentially bind to degenerate CANNTG E-box motifs to regulate gene expression (Borromeo et al., 2014; Borromeo et al., 2016; Casey et al., 2018; Castro et al., 2011). Using *de novo* motif analysis (Heinz et al., 2010), we identified the bHLH CAGCTG E-box motif as being highly enriched directly beneath 74% of the 13,457 ASCL1 combined peaks called, further confirming the quality of the ChIP-seq. Interestingly, we found that SOX and FOXO motifs were also significantly enriched within ASCL1 binding peaks (Figure 2C), suggesting that ASCL1 may function in combination with these transcription factor families to regulate gene expression in GBMs.

To identify putative-targets of ASCL1 in GBMs, we then used GREAT (McLean et al., 2010) to associate nearby genes that were upstream or downstream of the 13,457 ASCL1 binding peaks. From this analysis, we uncovered a total of 8,791 genes (red oval, Figure 2D). We reasoned that if these genes are regulated by ASCL1 then they should also be expressed in a manner correlated with *ASCL1* expression in GBMs. By applying Spearman’s rank-ordered correlation (>0.4) to RNA-seq of the 164 TCGA GBM samples, we then identified the top 10% of genes that showed a positive correlation with *ASCL1* expression across these tumor samples. We found 2,136 genes that are positively correlated with *ASCL1* expression (green oval, Figure 2D). When we cross referenced these 2,136 genes with the 8,791 genes identified from the ASCL1 ChIP-seq, there was an overlap of 1,106 genes, which we define as ASCL1 target genes in GBM (yellow area, Figure 2D). Supporting the validity of this approach, all ASCL1 canonical targets examined were included in this 1,106 putative-target gene list (Figure 2D, Table S2).

By evaluating the ASCL1 putative-target gene list, we uncovered a variety of genes that are particularly relevant to GBM development. Indeed, some of the most notable target genes include other neurodevelopmental and/or glial transcription factors such as OLIG genes (*OLIG1, OLIG2*), SOX genes (*SOX1, SOX2, SOX3, SOX4, SOX6, SOX8, SOX10*), NFI genes (*NFIA, NFIB, NFIX*), POU domain genes (*POU3F2, POU3F3, POU6F1*), Sal-like genes (*SALL2, SALL3*), and homeobox genes (*NKX2.2, ZEB1*). The functions of OLIG2 (Ligon et al., 2007; Lu et al., 2016; Mehta et al., 2011), SOX2 (Gangemi et al., 2009; Singh et al., 2017), and NFIA (Glasgow et al., 2017; Lee, Hoxha, & Song, 2017) have previously been reported to be important for regulating the tumorigenic property of glioma cell lines and in glioma mouse models. ASCL1 target genes also include numerous cell cycle (*CCND2, CCNE2, CDC25C, CDK4, CDK5R1, CSNK1E, E2F2, MCPH1, POLA2, PRIM2*), mitosis (*AURKB, BRSK1, MCM10, RCC2*), chromatin modification (*CHD3, CHD6, CHD7, KDM3B, KDM4B, SMARCA, SMARCB1, SMARCD1*), as well as oncogenic signal transduction related genes (*AKT3, EGFR, ERBB3, GSK3β, MYCN, PIK3R1, RTKN, TCF7L1, TCF7L2*) (Table S2). Strong ASCL1 binding peaks at the loci of some of these genes in the PDX-GBMs lines are illustrated (asterisks, Figure 2E-H).

We next wanted to know how the expression of the 1,106 ASCL1 putative-target genes sort across the various GBM subtypes using RNA-seq of the 164 primary GBMs. Heatmap and dendrogram analysis revealed that, similar to *ASCL1*, the 1,106 putative-target genes were highly expressed in the proneural and classical GBM subtypes, in the majority of neural and mixed GBM subtypes, but was mostly absent in the mesenchymal GBM subtype (Figure 2I). In all, 109 of the TCGA GBM samples were positive for the ASCL1 putative-target genes, while the remaining 55 samples express very little or low levels of the ASCL1 putative-targets.

To gain insights into the collective significance of the 1,106 ASCL1 putative targets, we then performed gene set over-representation analysis to annotate their function using ConsensusPathDB, a comprehensive collection of molecular interaction databases integrated from multiple public repositories (Herwig et al., 2016). Interestingly, the top most enriched pathway identified was cell cycle (Figure 2J). This is consistent with a previous report showing that positive and negative cell cycle regulators in neural progenitor cells are targets of ASCL1 (Castro et al., 2011). Other pathways that are also enriched for ASCL1 targets include those involved in chromatin segregation such as Aurora B Signaling and Amplification of Signal from Kinetochores, and intracellular signaling pathways such as those involved in PIP3 Activates AKT Signaling, Signaling by Rho GTPases, Hippo Signaling Pathway, and Wnt Signaling Pathway & Pluripotency. Finally, cancer pathways such as Retinoblastoma in Cancer, Hepatocellular Carcinoma, and Endometrial Cancer were also enriched for ASCL1 targets (Figure 2J, Table S3). Taken together, these findings suggest that ASCL1 is a transcriptional regulator at the epicenter of multiple biological processes that are fundamental to cancer development.

### ASCL1, OLIG2, and SOX2 are co-expressed in early and terminal stage tumors of a mouse glioma model

To functionally test ASCL1’s role in gliomagenesis *in vivo*, we began by characterizing the temporal expression pattern of ASCL1 along with OLIG2, SOX2, and glial lineage markers in brain tumors induced in a mouse model carrying floxed alleles of the tumor suppressor genes Neurofibromin 1 (*Nf1*) and tumor protein 53 (*Tp53*) (*Nf1*^*F/F*^*;Tp53*^*F/F*^) (Lin et al., 2004; Zhu et al., 2001). *NF1* and *TP53* are two of the most highly mutated genes in human GBM (2008; Brennan et al., 2013; Verhaak et al., 2010), and Cre-recombinase deletion of these two tumor suppressor genes (*Nf1;Tp53*^*CKO*^) in neural progenitors or glial precursor cells have previously been shown to be fully penetrant in producing glioma tumors in the brain of mice (Alcantara Llaguno et al., 2009; Alcantara Llaguno et al., 2015; Zhu et al., 2005). When mice carrying a *Glast*^*CreERT2/+*^ knock-in allele (Mori et al., 2006) was crossed with the *Rosa26-loxP-stop-loxP-tdTomato* (*R26R*^*LSL-tdTom*^) reporter line (Madisen et al., 2010), we found that tdTomato fluorescence was restricted in the brain of neonatal pups if tamoxifen was administered at E14.5 (Figure 3A-C), making *Glast*^*CreERT2/+*^ ideal to combine with the *Nf1*^*F/F*^*;Tp53*^*F/F*^ alleles to induce brain tumors.

**Figure 3.**
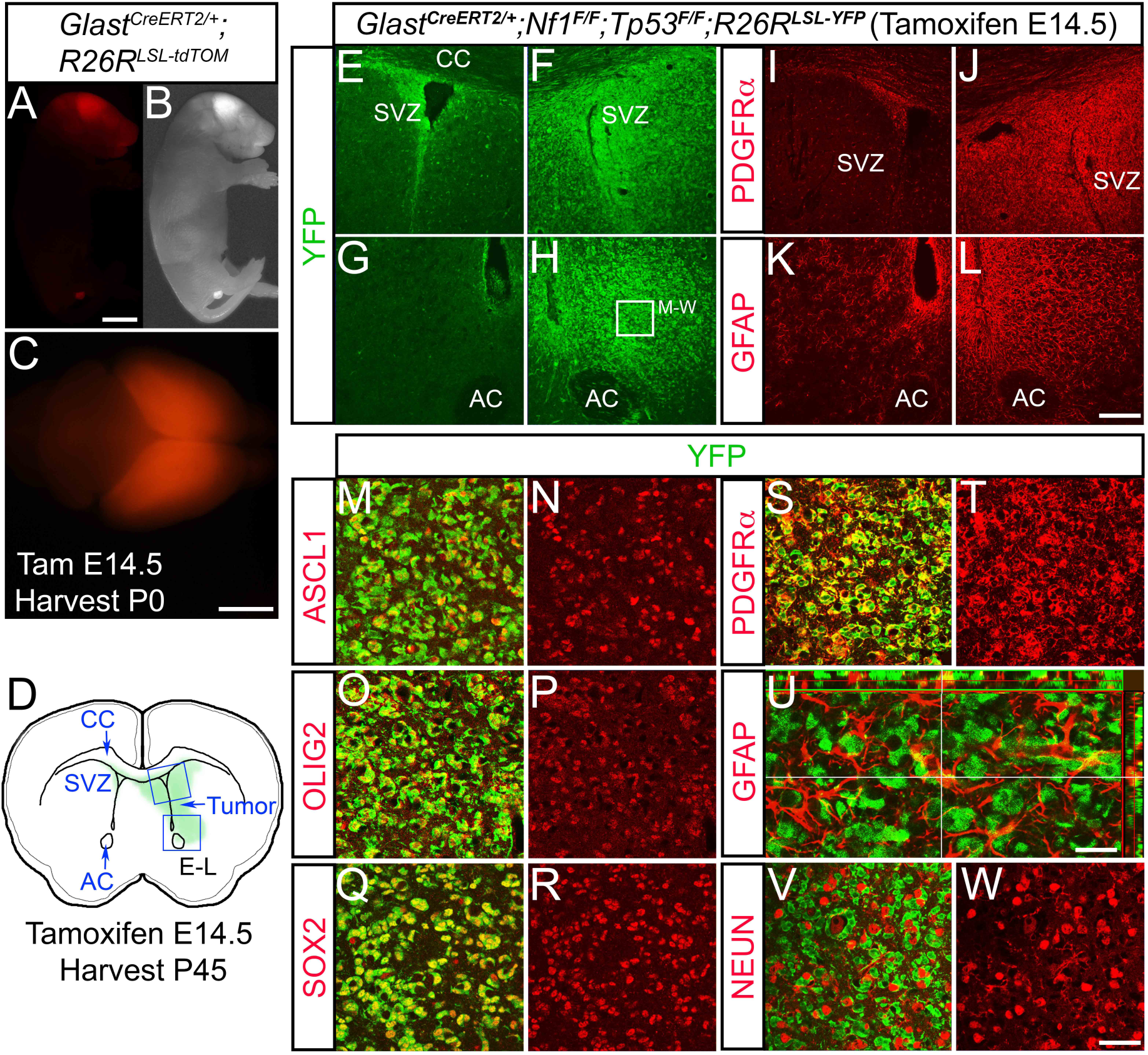
ASCL1, OLIG2, and SOX2 are highly expressed in early stage tumor cells of the glioma mouse model. **(A-C)** A neonatal pup from *Glast*^*CreERT2/+*^ crossed with *R26R*^*LSL-tdTomato*^ reporter administered with tamoxifen at E14.5. Note that tdTomato fluorescence is specific to the CNS and highest in the cerebral cortex (A,C). **(D)** Schematic of an early stage brain a tumor in the right subventricular zone (SVZ) of a *Glast*^*CreERT2/+*^*;Nf1*^*F/F*^;^*F/F*^*;R26R*^*LSL-YFP*^ mouse, administered with tamoxifen at E14.5 and harvested at P45. **(E-L)** Immunofluorescence shows high YFP reporter expression (E-H), OPC marker PDGFR*α* (I,J) and astrocyte marker GFAP (K,L) in tumor areas indicated in D. **(M-W)** Higher magnification of tumor area indicated in H showing ASCL1 (M,N), OLIG2 (O,P), SOX2 (Q,R), and PDGFR*α* (S,T) co-localized with YFP in tumor cells, but not GFAP (U) or the neuronal marker NEUN (V,W). Scale bar is 5 mm for A,B; 3 mm for C; 100 μm for E-L; 25 μm for M-T,V,W; and 12.5 μm for U.

To visualize the tumors as they develop in the brain, a *R26R*^*LSL-YFP*^ reporter allele (Srinivas et al., 2001) was incorporated into the glioma mouse model (*Glast*^*CreERT/+*^*;Nf1*^*F/F*^*;Tp53*^*F/F*^*;R26R*^*LSL-YFP*^). Tamoxifen was then administered to pregnant dams at E14.5 to induce *Nf1;Tp53*^*CKO*^ in neural progenitors of embryos. We first analyzed early tumors in the offspring at postnatal day (P) 45, at which point the majority of the mice were still asymptomatic and have yet to exhibit neurological symptoms. As expected, we were able to observe the presence of a tumor in some mice marked by intense YFP expression typically on one side of the brain surrounding the ventricle (Figure 3D-H). The tumor at this stage was not easily distinguishable from non-tumor tissues without YFP immunohistochemistry, yet both PDGFR*α*, an oligodendrocyte precursor cell (OPC) marker, and GFAP, an astrocyte marker, were ectopically expressed on the tumor side, indicating that the tumor is a glioma (Figure 3I-L). High magnifications revealed that ASCL1, OLIG2, and SOX2 are also expressed specifically within the YFP+ tumor cells (Figure 3M-R), and are highly irregular in shape, morphology, and density compared to normal YFP+ cells on the non-tumor side (not shown). Interestingly, the YFP+ tumor cells co-localized extensively with PDGFR*α* (Figure 3S,T) but not with GFAP or the neuronal marker NEUN (Figure 3U-W). The lack of co-localization between YFP and GFAP was similar to that observed in tumors of another glioma mouse model in which PDGF stimulation was combined with deletion of another tumor suppressor, *Pten* (Lei et al., 2011). This implies that the ectopic GFAP found infiltrating the YFP+ tumor tissue in our model may be reactive astrocytes rather than tumor cells themselves.

From P60-120, we found that 100% of *Nf1;Tp53*^*CKO*^ mice exhibited neurological symptoms and tumors that had evolved into an expanded mass with high mitotic index and microvascular proliferation resembling that of high grade gliomas (Figure 4A,B). We termed these *Ascl1*^*WT*^ tumor mice (N=29, blue line), which exhibited a median survival of 102 days, while CreER-negative littermate controls (N=19, green line) were tumor-free and healthy (Figure 4P). Over 90% of the tumors were found in the cortex and/or striatum area, while a minority were also found in olfactory bulb, diencephalon, midbrain, or cerebellum (Figure 4O). Similar to the early tumors and the PDX-GBMs, ASCL1, OLIG2, and SOX2 were co-expressed in the tumor cells of these terminal tumors, and many ASCL1+ tumor cells were also Ki67+ (Figure 4C-J). PDGFR*α* was also highly co-expressed by the ASCL1+ (Figure 4K) and OLIG2+ (not shown) tumor cells, whereas GFAP and to a lesser extent S100β and NEUN, although found in some parts of the tumor, did not overlap significantly with SOX2 or ASCL1 (Figure 4L-N).

**Figure 4.**
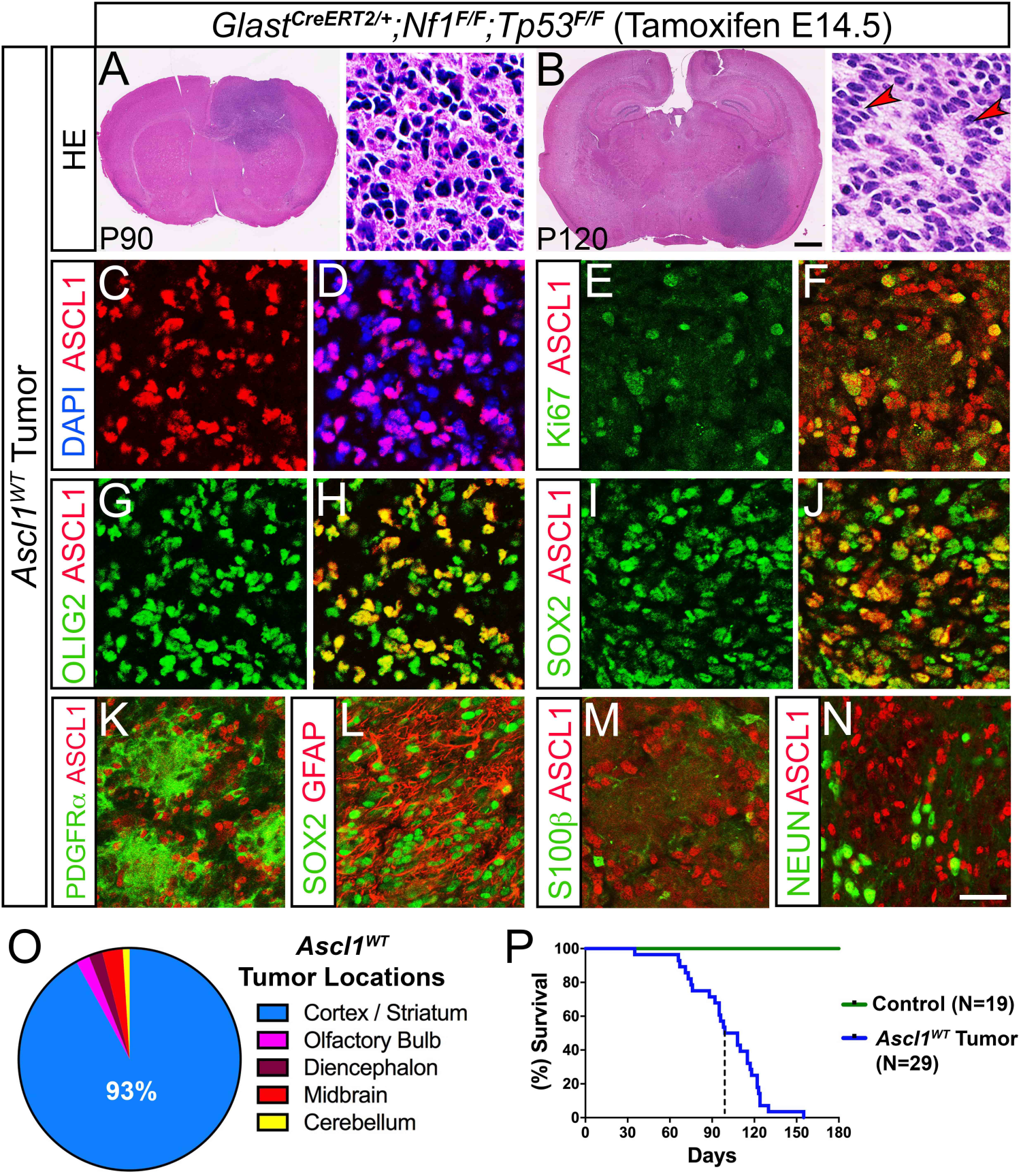
Expression of ASCL1, OLIG2, and SOX2 are maintained in mice with terminal stage glioma tumors. **(A,B)** H&E staining of *Ascl1*^*WT*^ terminal stage tumors harvested at P90 and P120. Higher magnification insets show that tumors are high-grade gliomas. Arrowheads indicate pseudopalisading cellular features consistent with GBM. **(C-N)** Immunofluorescence of *Ascl1*^*WT*^ GBM tumor tissue. ASCL1 is present in the majority of DAPI+ tumor cells (C,D) and co-localizes with Ki67 (E,F), OLIG2 (G,H), and SOX2 (I,J). PDGFR*α* (K) and GFAP (L) are also co-expressed in ASCL1+ or SOX2+ tumor cells respectively, but not S100β (M) and NEUN (N). **(O)** Incidence of tumors observed in different brain regions is indicated. Over 90% of tumors are found in the cortex and striatum (N=29). **(P)** Survival curve of *Ascl1*^*WT*^ tumor (N=29) bearing mice and Cre-negative control mice (N=19). Dotted line indicates median survival of 102 days for *Ascl1*^*WT*^ tumor mice. Scale bar is 1 mm for whole brain section and 30 μm for insets of A,B; and 25 μm for C-N.

Overall, our findings illustrate that ASCL1, OLIG2, and SOX2 are co-expressed in tumor cells of both early and terminal tumors of the glioma mouse model *in vivo*, and tumor cells maintain a molecular identity reminiscent of that of OPCs.

### Loss of ASCL1 decreases the proliferation of gliomas and increases the survival of tumor bearing mice

Currently, the direct requirement of ASCL1 in brain tumor formation and progression from low-grade gliomas to high-grade GBMs *in vivo* remains unknown. To address this, we incorporated *Ascl1*^*GFP*^ knock-in (null) and *Ascl1*^*Floxed*^ alleles into the glioma mouse model to generate *Glast*^*CreERT2*^*;Ascl1*^*GFP/F*^*;Nf1*^*F/F*^*;Tp53*^*F/F*^ and *Glast*^*CreERT2*^*;Ascl1*^*F/F*^*;Nf1*^*F/F*^*;Tp53*^*F/F*^ mice, respectively, both of which when administered with tamoxifen at E14.5 will result in triple conditional knock-out of *Ascl1* along with *Nf1* and *Tp53* (*Ascl1;Nf1;Tp53*^*CKO*^). To control for the possible effects of genetic background on glioma phenotype observed, we also generated *Glast*^*CreERT2*^*;Ascl1*^*GFP/+*^*;Nf1*^*F/F*^*;Tp53*^*F/F*^ and *Glast*^*CreERT2*^*;Ascl1*^*F/+*^*;Nf1*^*F/F*^*;Tp53*^*F/F*^ mice in parallel for comparison, both of which developed tumors that are still heterozygous for *Ascl1* when induced with tamoxifen, and are referred to as *Ascl1*^*HET*^ tumor mice.

Previous reports demonstrate that ASCL1 is essential for the proliferation of GBM cell lines *in vitro* (Park et al., 2017; Rheinbay et al., 2013). In contrast, *in vivo* we found that *Ascl1;Nf1;Tp53*^*CKO*^ mice (hence forth referred to as *Ascl1*^*CKO*^ tumor mice, N=39) still developed high-grade tumors that were phenotypically consistent with high grade gliomas (Figure 5A). Furthermore, *Ascl1*^*CKO*^ tumor penetrance and location (Figure 5L,M) in the brain was similar to the *Ascl1*^*HET*^ (not shown) and *Ascl1*^*WT*^ tumors (Figure 4O,P). We confirmed that ASCL1 was indeed absent in *Ascl1*^*CKO*^ tumors. As illustrated for a *Glast*^*CreERT2*^*;Ascl1*^*GFP/F*^*;Nf1*^*F/F*^*;Tp53*^*F/F*^ mouse, GFP driven by the endogenous *Ascl1* locus marks precisely the tumor cells but ASCL1 was no longer detected (Figure 5B,C). Notably, OLIG2 and SOX2 (Figure 5D,E,J,K), which we identified as ASCL1 target genes, were still expressed, indicating that expression of these two transcription factors do not depend solely on ASCL1. Similarly, OPC markers such as PDGFR□, the chondroitin sulfate NG2, and SOX10 were still expressed in the *Ascl1*^*CKO*^ tumors (Figure 5D,F,G). As observed in *Ascl1*^*WT*^ tumors, GFAP did not co-colocalize extensively with GFP+ tumor cells, despite being expressed in some regions of the tumor (Figure 5H,I). Together, these findings demonstrate that glial transcription factors and the OPC-like identity of the tumor cells are still retained in the absence of ASCL1.

**Figure 5.**
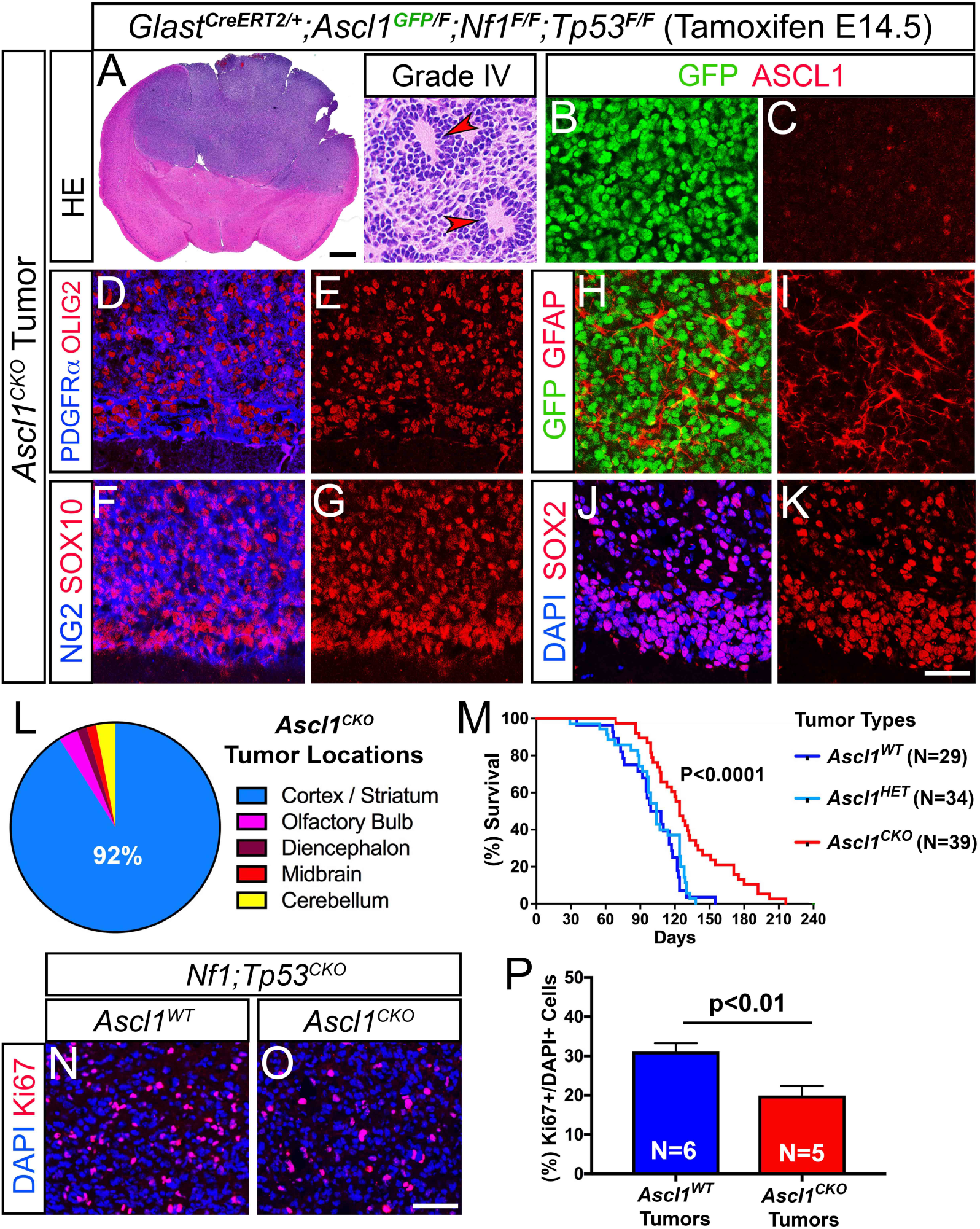
Survival of glioma tumor bearing mice is increased in the absence of ASCL1. **(A)** H&E staining of an *Ascl1*^*CKO*^ tumor exhibiting pseudopalisading cellular features of Grade IV glioma (arrowheads, insets). **(B-K)** Immunofluorescence of *Ascl1*^*CKO*^ tumor. GFP, driven by an *Ascl1*^*GFP*^ knock-in allele, is present in tumor cells but ASCL1 is absent (B,C), indicating efficient deletion of *Ascl1*^*Floxed*^ allele. Expression of OLIG2 & PDGFR*α* (D,E), SOX10 & NG2 (F,G), GFAP (H,I) and SOX2 (J,K) are unaffected. **(L)** Incidence of *Ascl1*^*CKO*^ tumors observed in the different brain regions. Over 90% of tumors are found in the cortex and striatum area similar to *Ascl1*^*WT*^ tumors. **(M)** Survival curve of *Ascl1*^*CKO*^ versus *Ascl1*^*HET*^ tumor mice. Median survival is significantly improved for *Ascl1*^*CKO*^ (122 days) compared to *Ascl1*^*HET*^ (104 days) tumor mice (dotted lines). Note that survival of *Ascl1*^*HET*^ is very similar to *Ascl1*^*WT*^ tumor mice (see Fig. 5P). **(N-P)** Immunofluorescence (N,O) and quantification of the percentage of Ki67+/DAPI+ tumor cells (P) for *Ascl1*^*WT*^ and *Ascl1*^*CKO*^ tumors. Scale bar is 1 mm for whole brain section and 30 μm for insets of A; 25 μm for B-K; and 50 μm for N,O.

Notably, the *Ascl1*^*CKO*^ tumor mice survived longer compared to *Ascl1*^*HET*^ and *Ascl1*^*W*T^ tumor mice. Specifically, while the *Ascl1*^*HET*^ tumor mice (N=34) died between P60-130, with a median survival of 104 days, which is very similar to *Ascl1*^*WT*^ tumor mice (median survival of 102 days), *Ascl1*^*CKO*^ tumor mice (N=39) died later between P90-180, with a median survival of around 122 days (compare red versus light and dark blue lines, Figure 5M). This improvement in survival for the *Ascl1*^*CKO*^ tumor mice also holds true even when analyzed by gender (not shown) and strongly suggests that it was due to the loss of ASCL1.

To determine what may account for the improved survival of the *Ascl1*^*CKO*^ tumor mice, we assessed tumor proliferation by quantifying the percentage of tumor cells that were Ki67+ in comparison to *Ascl1*^*WT*^ tumor mice. Because the density of Ki67+ cells can vary dramatically across a large tumor depending on necrosis or the integrity/quality of the tumor tissue, we chose to image and quantify several regions of each *Ascl1*^*CKO*^ (N=5) or *Ascl1*^*WT*^ tumor (N=6) with the highest density of Ki67+ cells (Figure 5N,O). Overall, *Ascl1*^*CKO*^ tumors exhibited a decrease of about 30% Ki67+ cells compared to *Ascl1*^*WT*^ tumors (Figure 5P), which is consistent with our previous finding that numerous cell cycle and mitotic genes are targets of ASCL1. This decrease in Ki67+ cells was similar to that observed for adult OPCs in the spinal cord when *Ascl1* was conditionally deleted (Kelenis, Hart, Edwards-Fligner, Johnson, & Vue, 2018) and supports the interpretation that the increased survival of *Ascl1*^*CKO*^ tumor mice may result from a decrease in the rate of tumor cell proliferation.

### Transcriptome of mouse GBM tumors showed that loss of ASCL1 is associated with down-regulation of cell cycle genes

To determine if the loss of ASCL1 altered the molecular profiles of the mouse glioma tumors, we carefully isolated tumor tissues from *Ascl1*^*WT*^ (N=5) and *Ascl1*^*CKO*^ (N=5) tumor mice for bulk RNA-seq analysis. RNA-seq tracks of the *Ascl1* locus show that *Exon 1* and *2* of the *Ascl1 mRNA* (containing the entire coding sequence), were completely absent in all *Ascl1*^*CKO*^ tumors but were present in *Ascl1*^*WT*^ tumors (Figure 6A), confirming efficient deletion of the *Ascl1*^*Floxed*^ allele. We first compared the transcriptomes of the *Ascl1*^*WT*^ and *Ascl1*^*CKO*^ tumors with transcriptomes of CNS cell types, including OPCs, newly formed oligodendrocytes (NFO), mature oligodendrocytes (MO), astrocytes (AS), neurons, and whole cortex (WC) (Zhang et al., 2014). A multidimensional scaling (MDS) plot shows that both the *Ascl1*^*WT*^ and *Ascl1*^*CKO*^ tumors cluster together and away from the CNS cell types, and therefore are more similar to each other than to neurons or any of the glial lineage cells (Figure 6B). When we further analyzed RNA-seq of *Ascl1*^*WT*^ and *Ascl1*^*CKO*^ tumors using the top 50 signature genes for each CNS cell type, both tumor types more closely resemble that of OPCs versus the other CNS cell types (Figure 6C). This finding further supports the notion that OPCs, which are highly proliferative, may be the precursor cell-of-origin for the glioma tumors in this model.

**Figure 6.**
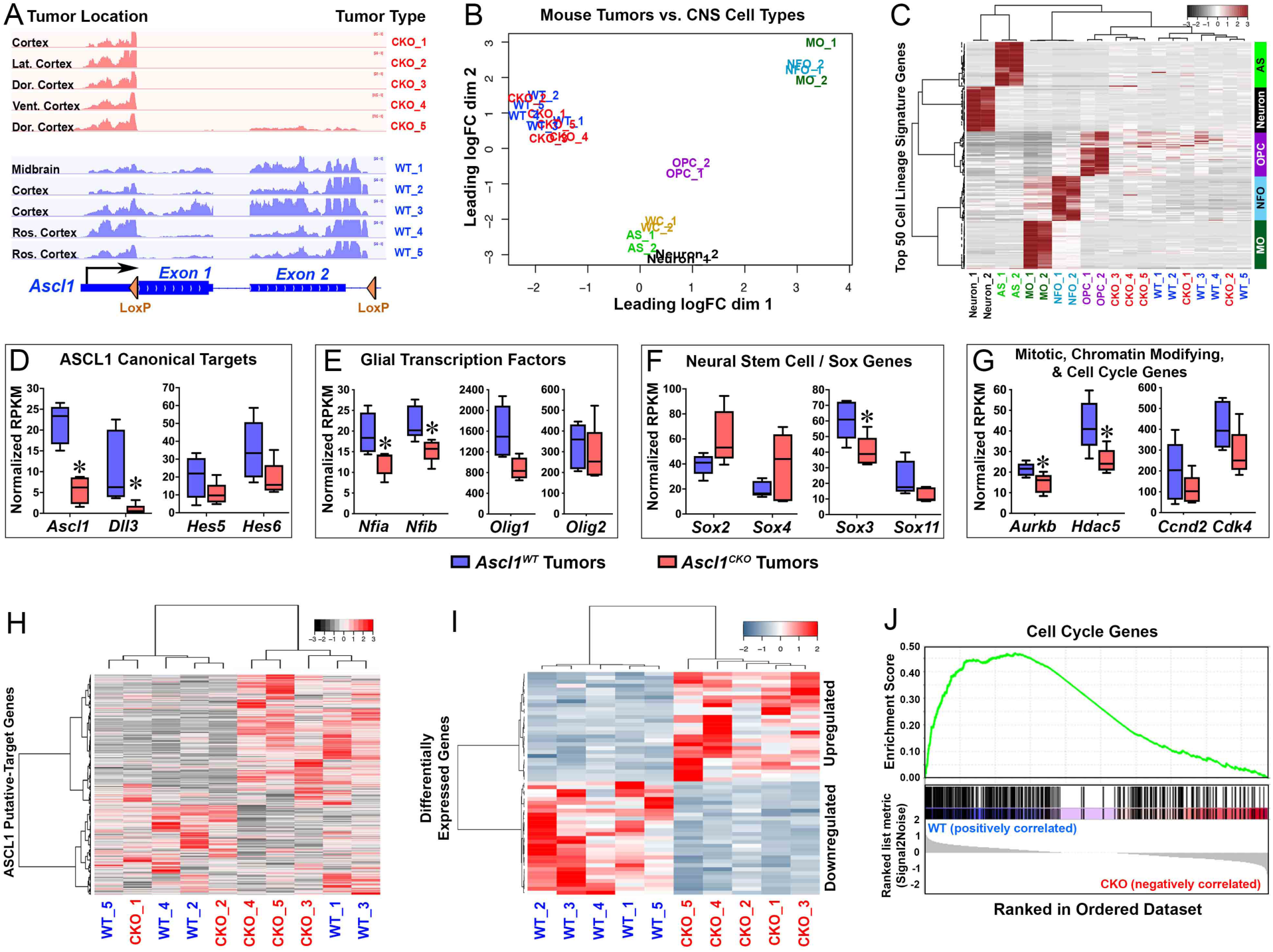
Cell cycle genes are down-regulated in *Ascl1*^*CKO*^ glioma tumors of the mouse model. **(A)** RNA-seq tracks at the *Ascl1* locus of *Ascl1*^*WT*^ and *Ascl1*^*CKO*^ tumors isolated from brain regions indicated. Note that Exon 1 and 2 of the *Ascl1 mRNA*, flanked by Lox P sites, are absent in *Ascl1*^*CKO*^ tumors. **(B)** Multidimensional scaling (MDS) plot of RNA-seq of *Ascl1*^*WT*^ and *Ascl1*^*CKO*^ tumors versus CNS cell types (Zhang et al, 2014). *Ascl1*^*WT*^ and *Ascl1*^*CKO*^ tumors are more similar to each other than to any of the CNS cell types. AS – astrocytes; OPC – oligodendrocyte precursor cells; NFO – newly formed oligodendrocytes; MO – myelinating oligodendrocytes; WC – whole cortex. **(C)** Heatmap and dendrograms using the top 50 CNS cell lineage signature genes for each cell type (Zhang et al, 2014). Dendrograms on top show that *Ascl1*^*WT*^ and *Ascl1*^*CKO*^ tumors express signature genes that are more similar to OPCs than to the other CNS cell types. **(D-G)** Box and whisker plots of ASCL1 putative-target genes in *Ascl1*^*WT*^ and *Ascl1*^*CKO*^ tumors. Canonical targets of ASCL1 (E), glial transcription factors (G), and mitotic, chromatin modifying, and cell cycle genes (H) are expressed at lower level while *Sox* genes (F) are bidirectionally affected in *Ascl1*^*CKO*^ compared to *Ascl1*^*WT*^ tumors. Asterisks indicate target genes significantly altered (P<0.05, Wilcox test). **(H,I)** Heatmap and dendrograms of differentially expressed genes (DEGs) in *Ascl1*^*WT*^ and *Ascl1*^*CKO*^ GBMs. ASCL1 putative direct targets that are upregulated or downregulated (I, Supplementary Table S5) and 57 ASCL1 indirect target DEGs (FDR<0.05) were identified (J, Supplementary Table S6). **(J)** Gene-set-enrichment-analysis (GSEA) showing that cell cycle genes are enriched in the downregulated genes in *Ascl1*^*CKO*^ compared to *Ascl1*^*WT*^ tumors.

Finally, we sought to identify genes that are differentially expressed between *Ascl1*^*WT*^ and *Ascl1*^*CKO*^ tumors. Analysis of ASCL1 canonical target genes revealed that *Dll3*, similar to *Ascl1*, was significantly decreased, while *Hes5* and *Hes6* were lowered (Figure 6D), but *Dll1, Notch1*, and *Insm1* (not shown) were unchanged in *Ascl1*^*CKO*^ tumors. Interestingly, glial transcription factors *Nfia* and *Nfib*, and several mitotic (*Aurkb*) and chromatin modifying (*Hdac5*) genes were significantly decreased, whereas *Olig1, Olig2*, and cell cycle genes (*Ccnd2*, and *Cdk4*) were modestly reduced in *Ascl1*^*CKO*^ tumors (Figure 6E,G). In contrast, *Sox* genes were bidirectionally affected by the loss of ASCL1. For instance, although *Sox3* and *Sox11* were decreased, *Sox2* and *Sox4* appeared upregulated in the *Ascl1*^*CKO*^ tumors (Figure 6F). Heatmap and dendrogram analysis of all 1,054 ASCL1 putative targets (converted from a list of 1,106 genes from human GBMs, Table S3), revealed that there were as many genes being upregulated as there were genes being downregulated by the loss of ASCL1 (Figure 6H, Table S4). We also identified over 50 indirect targets of ASCL1 that were either upregulated or downregulated in the *Ascl1*^*CKO*^ tumors (Figure 6I, Table S5). Finally, in agreement with our earlier finding that tumor cell proliferation is decreased in the absence of ASCL1, gene-set-enrichment analysis revealed that cell cycle related genes were highly enriched in the down-regulated genes in the *Ascl1*^*CKO*^ tumors (Figure 6J). This suggests that a decreased in cell-cycle related gene expression may contribute to the increase in survival of the *Ascl1*^*CKO*^ tumor mice.

In summary, our findings highlight an *in vivo* role for ASCL1 in modulating the expression of a variety of genes, including neurodevelopmental or glial transcription factors and cell cycle genes, either directly or indirectly, that are crucial for the proliferation of GBM tumors in the brain.

## DISCUSSION

We demonstrate in this study that ASCL1 is highly expressed in the majority of PDX-GBM cells *in vivo*, with over 90% of ASCL1+ cells co-expressing OLIG2 and SOX2. Interestingly, in addition to OLIG2 and SOX2, we find that expression of a variety of other genes encoding transcription factors such as NFI, POU domain, Sal-like, SOX, as well as homeobox are also highly correlated with *ASCL1* expression in RNA-seq of primary GBMs (Table S2). This finding is similar to that previously reported in GSCs from cultured GBM cell lines (Rheinbay et al., 2013; Suva et al., 2014). Accordingly, we find that these transcription factor encoding genes are not only correlated with *ASCL1* expression but are also targets of ASCL1 binding (Table S1). These findings support a complex transcription factor interaction network in which the co-expression of these transcription factors may be interdependent on each other, and this co-expression is essential for regulating genes crucial for maintaining GSCs in an aberrant stem-like state of dedifferentiation and proliferation. In agreement with this, it is not surprising that combinatorial over-expression of multiple transcription factors is necessary and sufficient to reprogram differentiated glioma cells or immortalized astrocytes into tumor propagating cells (Singh et al., 2017; Suva et al., 2014).

ChIP-seq for ASCL1 has previously been performed for GBM cell lines in culture revealing that ASCL1 directly interacts with Wnt signaling by binding to genes such as *AXIN2, DKK1, FZD5, LGR5, LRP5, TCF7* and *TCF7L1*. A model was proposed in which ASCL1 functions at least in part by repressing an inhibitor of Wnt signaling, *DKK1*, resulting in increased signaling through this pathway to maintain the tumorigenicity of GBM cells (Rheinbay et al., 2013). Similarly, in this study we find that all of the aforementioned genes as well as numerous other Wnt related genes (*GSK3B, LRP4, LRP6, TCF7L2*) are directly bound by ASCL1 (Table S1). Additionally, expression of many of these Wnt related genes is positively correlated with *ASCL1* expression when analyzed across RNA-seq of the 164 TCGA primary GBM samples, and Wnt Signaling and Pluripotency was identified as one of the pathways significantly over-represented by the ASCL1 target genes that we identified in this study (Figure 2J, Table S2). Despite these findings, RNA-seq from mouse *Ascl1*^*CKO*^ tumors revealed that expression of many of the Wnt related genes was unaffected by the loss of ASCL1. Thus, although ASCL1 binds to and may contribute to the regulation of some of these genes, particularly in an *in vitro* setting (Rheinbay et al., 2013), expression of Wnt pathway genes remains in GBMs *in vivo* in the absence of ASCL1. Consequently, the presence of Wnt signaling may partly contribute to the formation of *Ascl1*^*CKO*^ tumors in the brain of the glioma mouse model.

In addition to gliomas, ASCL1 is highly expressed in cancers with neuroendocrine characteristics from multiple tissues including small cell lung carcinoma (SCLC), prostate cancer, and thyroid medullary carcinoma (Chen, Kunnimalaiyaan, & Van Gompel, 2005; Rapa et al., 2013; Zhang et al., 2018). Previously, we reported that ASCL1 is required for tumor formation in a mouse model of SCLC (Borromeo et al., 2016). This finding reflects the requirement for ASCL1 in the generation and survival of pulmonary neuroendocrine cells (PNECs), a presumptive cell-of-origin for SCLC. In contrast, in this study we found that ASCL1 is not required for GBM formation in the brain of the glioma mouse model, although disease progression is altered and the animals have extended survival. Based on cell lineage markers in the glioma mouse model used here, OPCs are implicated as the presumptive cell-of-origin for the tumors. OPCs are known for displaying highly migratory and proliferative behavior similar to GBM. OPC specification and generation in the CNS is dependent upon OLIG2 (Lu et al., 2002; Zhou, Choi, & Anderson, 2001), however ASCL1 also plays an important role to regulate the number and proliferation of OPCs (Kelenis et al., 2018; Nakatani et al., 2013; Parras et al., 2007; Vue et al., 2014). Interestingly, in addition to ASCL1 and OLIG2, transcription factors such as NFIA, SOX2, and SOX10 are also expressed in OPCs. However, as OPCs differentiate to become mature oligodendrocytes, only OLIG2 and SOX10 are maintained while ASCL1, NFIA, and SOX2 are down-regulated (Glasgow et al., 2014; Laug, Glasgow, & Deneen, 2018; Nakatani et al., 2013). This down-regulation suggests that the co-expression of these transcription factors is important for maintaining OPCs in a progenitor-like state, and the loss of just one of these factors does not completely abrogate tumor formation following deletion of *Nf1* and *Tp53* because OPCs are still generated, and are thus susceptible to being transformed into glioma. The direct roles of NFIA and OLIG2 in tumor development in glioma mouse models were also previously tested. Similar to the findings for ASCL1, tumor formation persisted in the absence of each of these transcription factors. Furthermore, despite utilizing different approaches and driver mutations to induce tumor formation, the loss of NFIA or OLIG2 was also accompanied by significant decreases in tumor cell proliferation resulting in an increase in survival for their respective mouse models (Glasgow et al., 2017; Lu et al., 2016). Together, these studies illustrate potential redundant roles for neurodevelopmental or glial transcription factors in driving GBM formation and progression *in vivo* in the brain, where the loss of one factor is likely to be compensated by the remaining transcription factors.

Similar to our study here, deletion of *Nf1, Tp53*, along with or without *Pten*, was previously shown to be fully penetrant in producing GBM tumors in the brain of mice (Alcantara Llaguno et al., 2009; Alcantara Llaguno et al., 2015; Zhu et al., 2005). More specifically, tumors were successfully induced from neural stem cells in the SVZ of the lateral ventricles, including ASCL1+ transiently amplifying progenitors, as well as OPCs. It was reported that two types of glioma tumors were observed when *Nf1* and *Tp53* were deleted in the adult brain (Alcantara Llaguno et al., 2015). Type 1 tumors, which are found in dorsal/anterior brain regions such as striatum, hippocampus, and cortex, are highly infiltrative and aggressive. Type 2 tumors, on the other hand, are found more in ventral/posterior brain regions such as the diencephalon and brainstem, and exhibit well-defined boundaries. Based on gene expression, Type 1 tumors express high levels of GFAP and are speculated to be derived from neural stem cells in the SVZ, whereas Type 2 tumors express high levels of OLIG2 and PDGFR□, and likely to be derived from OPCs. Over 90% of the mouse GBM tumors that we observed in this study are found in the telencephalon, predominantly in the cortex and striatum, suggesting that they may be similar to Type 1 tumors. However, although GFAP expression is high in some of the tumors of our model, the majority of GFAP expressing cells do not seem to co-localize with Cre-reporters such as YFP, which directly mark the tumor cells. Instead, YFP colocalizes extensively with OLIG2 and PDGFR□, indicating that they are more similar to Type 2 tumors in terms of gene expression. This inconsistency likely reflects our use of *Glast*^*CreERT2*^ to delete *Nf1* and *Tp53*, which may target neural progenitors in the SVZ as well as glial precursor cells outside of the SVZ, and the timing of tumor induction embryonically in our model rather than in the adult brain.

In conclusion, the tumors induced in our mouse model are highly heterogenous based on RNA-seq analysis, which is similar to that seen for human GBMs (Patel et al., 2014; Sottoriva et al., 2013). This heterogeneity is likely the result of different tumors being spontaneously derived from different cell-of-origins in the various brain regions, and are thus exposed to different microenvironments. Despite this heterogeneity, however, the loss of ASCL1 still significantly delays tumor progression and resulted in a significant increase in survival for *Ascl1*^*CKO*^ tumor mice, illustrating an important role for ASCL1 in controlling the rate of GBM proliferation *in vivo*. A fundamental question remaining for future studies is whether ASCL1 and other transcription factors are similarly required directly within growing GBMs in the brain, and how much these transcription factors may contribute to GBM recurrence, if any, following multimodal treatments.

## ACKNOWLEDGEMENTS

We acknowledge Lauren K. Tyra, Erin Kibodeaux, Juan Villareal, and Trisha Savage for excellent mouse genotyping and husbandry. We thank Dr. Francois Guillemot for *Ascl1*^*flox*^ mice and Dr. Luis F. Parada for *Nf1*^*flox*^*;Trp53*^*flox*^ mice, the Histo Pathology Core for performing H&E staining of brain tumor tissues, and members of the J.E.J laboratory for helpful discussions throughout this study. This research was supported by the National Institutes of Health (NIH) R01 NS032817 and CPRIT RP130464 to J.E.J, and NIH F32 CA168330 and K22 NS092767 to T.Y.V. This research was also partially supported by UNM Comprehensive Cancer Center Support Grant NCI P30CA118100 and made use of the Fluorescence Microscopy and Cell Imaging shared resource.

## SUPPLEMENTARY FIGURES & TABLES

**Figure S1.**
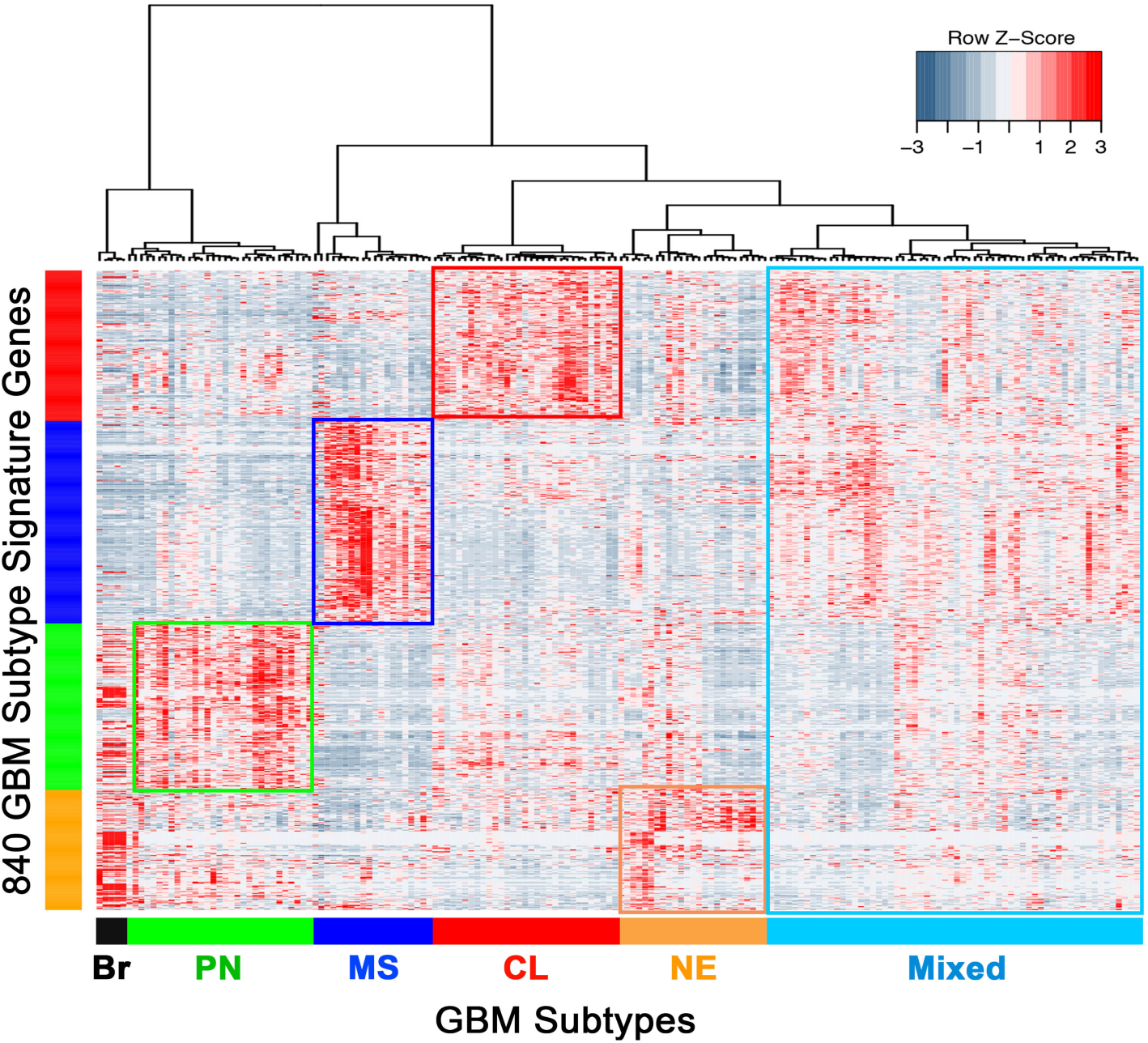
Subtype identities of primary GBMs using RNA-seq. RNA-seq data of 164 TCGA Primary GBMs and 5 normal brain samples (Brennan et al, 2013). Heatmap and dendrogram using the 840 GBM Subtype Signature Genes (Verhaak et al, 2010) reveals the presence (rectangles) of four previously identified GBM subtypes (PN-proneural, MS-mesenchymal, CL-classical, NE-neural) as well as Mixed GBM group which express multiple subtype signatures (P).

**Table S1: ChIP-seq coordinates of ASCL1 binding sites in PDX-GBMs**

**Table S2: ASCL1 putative target genes (Venn Diagrams in Figure 2D)**

**Table S3: ConsensusPathDB Analysis: Complete list of gene sets enriched in ASCL1 putative target genes (include biological pathway terms and the genes in each selected pathways) (Diagram in Figure 2J)**

**Table S4: Expression of ASCL1 putative target genes in mouse *Ascl1***^***WT***^ **and *Ascl1***^***CKO***^ **tumors (Figure 6H)**

**Table S6: List of differentially expressed genes in mouse *Ascl1***^***WT***^ **and *Ascl1***^***CKO***^ **tumors (Figure 6I)**

